# Establishing RPTE-derived cell lines expressing hTERT for studying BK polyomavirus

**DOI:** 10.1101/766675

**Authors:** Linbo Zhao, Michael J. Imperiale

**Affiliations:** Department of Microbiology and Immunology, University of Michigan, Ann Arbor, MI, 48109, USA; Comprehensive Cancer Center, University of Michigan, Ann Arbor, MI, 48109, USA

**Author notes:** Corresponding author. Department of Microbiology and Immunology, 1150 West Medical Center Drive, 5724 Medical Science II, Ann Arbor, MI 48109-5620, USA. E-mail addresses (L. Zhao); (M.J. Imperiale).

## Abstract

We previously established an infection model for BKPyV in primary human renal proximal tubule epithelial (RPTE) cells. Use of these cells is limited by their inability to be passaged extensively. We describe RPTE cells immortalized with hTERT, which can serve as a model system for acute or persistent BKPyV infection.

## ANNOUNCEMENT

BK Polyomavirus (BKPyV) was first isolated in 1971 (1). BKPyV is a member of the *Polyomaviridae*, which is a group of small, icosahedral, nonenveloped viruses with circular double-stranded DNA genomes. Several systems for studying BKPyV rearranged variants, which are generally isolated from patients with BKPyV disease and can replicate robustly in culture, have been established (2, 3). Our lab previously demonstrated that primary human renal proximal tubule epithelial (RPTE) cells could serve as an acute lytic infection model for studying the BKPyV Dunlop variant, and other variants, in vitro (4). One of the disadvantages of primary RPTE cells is that they divide slowly and stop growing at around passage 10. To extend the time window for BKPyV research, we generated human telomerase reverse transcriptase (hTERT)-expressing RPTE (RPTE-hTERT) cells.

RPTE cells were acquired from Lonza and grown according to our previous report (5). RPTE and RPTE-hTERT cells were maintained in Renal Epithelial Cell Growth Medium Kit (REGM/REBM, Lonza). An earlier publication showed that hTERT can be used to immortalize RPTE cells (6). However, the lentivirus vector (pLXSN) used in the previous report contains an SV40 origin of replication, which could lead to problems due to its ability to replicate in the presence of BKPyV. We therefore started with another lentivirus vector to avoid the use of SV40 sequences. pLenti CMV GFP Puro (658-5) was a gift from Eric Campeau & Paul Kaufman (Addgene plasmid #17448) (7). hTERT was amplified with primer pairs (XbaI-Kozak-hTERT-F, 5’ AAATCTAGAGCCGCCACCATGCCGCGCGCTCCCCGCTGC 3’ and SalI-hTERT-R, 5’ AGGGTCGACTCAGTCCAGGATGGTCTTGAA 3’). We first prepared an intermediate plasmid (pLenti-CMV-hTERT-puro) by substituting GFP in the pLenti CMV GFP Puro plasmid with hTERT at the XbaI and SalI sites (Figure 1A). In the second step, the WPRE sequence was amplified with primer pairs (SalI-WPRE-F, 5’ TGAGTCGACAATCAACCTCTGGAT 3’ and KpnI-WPRE-R, 5’ AAAGGTACCAGGCGGGGAGGCGGCCCAA 3’), and the puromycin selection markers in the pLenti-CMV-hTERT-puro were deleted to construct the final pLenti-CMV-hTERT plasmid by substituting the fragment between KpnI and SalI sites with the amplified WPRE. The hTERT-WPRE region was sequenced to confirm the integrity of PCR amplification and cloning.

**Figure 1.**
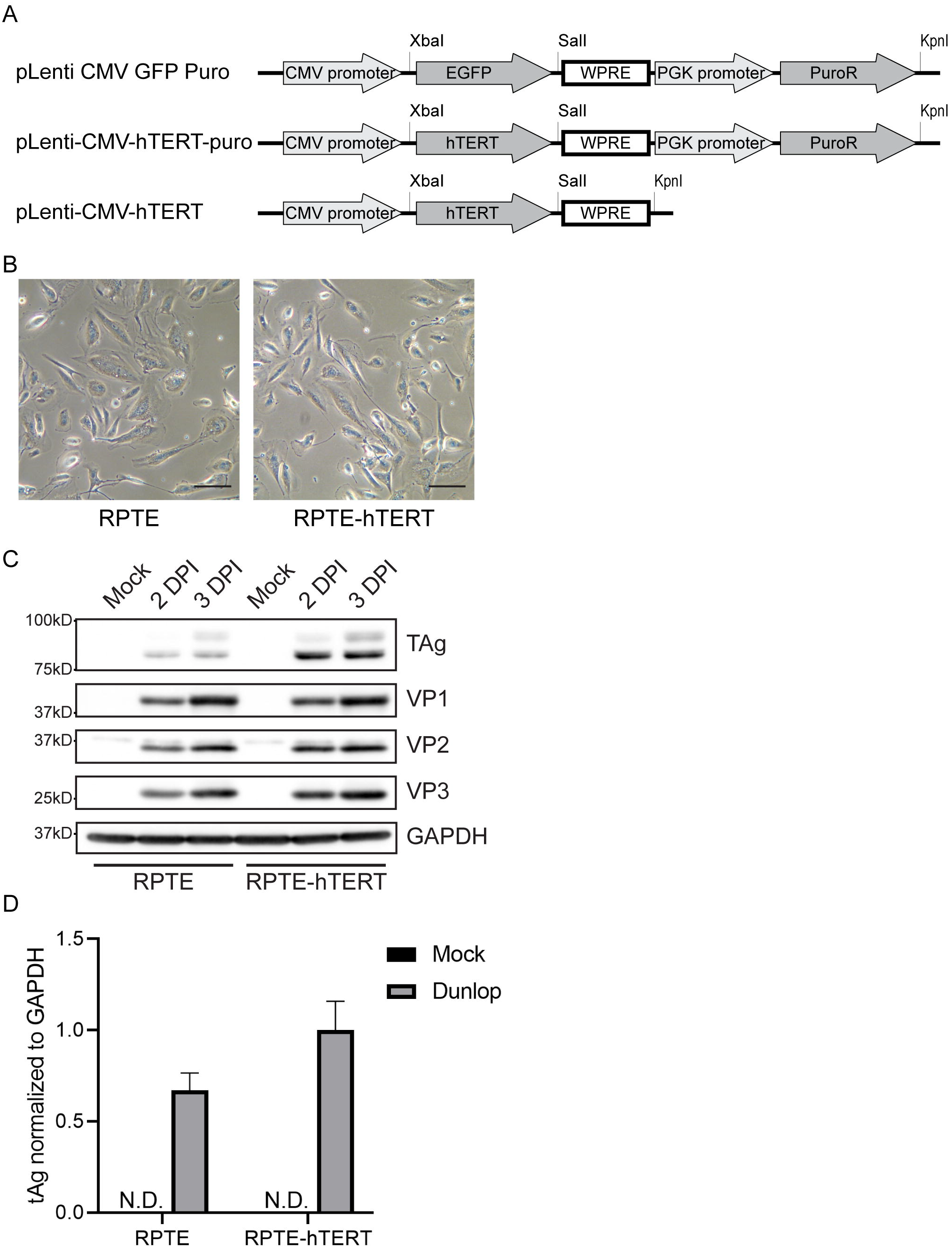
A: Schematic diagram of the lenvirus plasmid construction. B: Phase-contrast images of RPTE and RPTE-hTERT cells. RPTE cells were plated one day before taking images. Images were taken using a phase-contrast microscope. Bars represent 100 μm. C: Viral protein expression in RPTE and RPTE-hTERT cells. RPTE or RPTE-hTERT cells were infected by BKPyV at a MOI of 1. Viral early protein large tumor antigen (TAg); late proteins VP1, VP2, and VP3; and GAPDH were examined by Western Blot. D: Viral early protein small tumor antigen (tAg) expression in RPTE and RPTE-hTERT cells. RPTE or RPTE-hTERT cells were infected by BKPyV at a MOI of 1. tAg mRNA expression was examined by RT-qPCR.

hTERT-expressing lentivirus was produced by co-transfecting pLenti-CMV-hTERT, pRSV-Rev, pMDLg/pRRE, and pMD2.G plasmids into 293TT cells (8, 9). Fresh media was supplied 16 hours post-transfection. Media containing lentivirus was harvested at 48- and 72-hours post-transfection and filtered with a 0.45 μm polyethersulfone filter (MilliporeSigma). Filtered media was overlaid on 20% sucrose in 1x phosphate buffered saline, and lentivirus was concentrated by centrifuging at 24,000 rpm for 2 hours (AH-629 rotor). Pelleted lentiviruses were resuspended in complete REGM/REBM.

RPTE cells at passage 3 were grown in REGM/REBM media in a 10 cm dish. hTERT-expressing lentivirus at a multiplicity of infection (MOI) of 0.3 was directly added to the cells and inoculated at 37 °C overnight. Cells were passaged at 3 days post-transduction and further passaged at 70% confluency until passage 20 to select against non-transduced cells. Single RPTE-hTERT subclones were isolated by seeding hTERT-transduced cells at passage 20 in 96 well plates at a concentration of 0.2 cells per well. Subclones were subsequently expanded in 6 well plates and 10 cm dishes before freezing down aliquots in REBM/REGM with 10% DMSO and 10% FBS in liquid nitrogen. hTERT integration was confirmed by amplifying hTERT-WPRE fragment from cellular genomic DNA and Sanger sequencing (data not shown). Images of RPTE-hTERT cells are shown in Figure 1B.

To test if RPTE-hTERT cells are susceptible to BKPyV infection, RPTE-hTERT cells and RPTE cells were infected with BKPyV (Dunlop) at a MOI of 1 as previous described (10). Protein samples were harvested with E1A buffer (50 mM HEPES [pH 7], 250 mM NaCl, and 0.1% NP-40, with inhibitors: 5 μg/ml PMSF, 5 μg/ml aprotinin, 5 μg/ml leupeptin, 50 mM sodium fluoride and 0.2 mM sodium orthovanadate added right before use). Equal protein was electrophoresed on a 4-15% precast protein gel (Bio-Rad). The separated proteins were transferred to nitrocellulose membranes with the Trans-Blot Turbo Transfer System (Bio-Rad). Membranes were blocked in 2% nonfat milk in PBS-T buffer (144 mg/L KH_2_PO_4_, 9 g/L NaCl, 795 mg/L Na_2_HPO_4_, pH 7.4, 0.1% Tween 20) for 1 hour at room temperature. Antibodies for Western blotting were diluted in 2% milk in PBS-T as follows: anti-large tumor antigen (TAg) mouse ascites (pAb416) at 1:5,000 (11); anti-glyceraldehyde 3-phosphate dehydrogenase (GAPDH, MilliporeSigma, CB1001) at 1:200,000; anti-VP1 (pAb5G6) mouse ascites at 1:5000; custom made rabbit anti-VP2 (Bethyl Labs) at 1:10,000 (12); horseradish peroxidase (HRP)-conjugated ECL sheep anti-mouse (GE healthcare, NA931V) at 1: 5,000; and HRP-conjugated ECL donkey anti-rabbit antibody (GE Healthcare, NA934V) at 1: 5,000. The probed membrane was developed with HRP substrate (MilliporeSigma, WBLUF0100), and images were acquired with the Syngene PXi gel doc system. Western blotting showed that there was a slight increase of viral early protein TAg at 48 hours post-infection in the RPTE-hTERT cells as compared to the RPTE cells (Figure 1C), while late protein VP1, VP2, and VP3 expression is similar between RPTE and RPTE-hTERT cells, which suggests an early accelerated early phase of the virus life cycle because the RPTE-hTERT cells are actively dividing. Due to lack of small tumor antigen (tAg) antibody, tAg expression was examined by reverse transcription-quantitative PCR (RT-qPCR). Total cellular RNA was harvested with TRIzol RNA isolation reagent (Thermo Fisher Scientific) and purified with a spin column-based purification kit (Zymo Research). cDNA was synthesized with SuperScript III Reverse Transcriptase (Thermo Fisher Scientific). qPCR was performed on the cDNA with PowerUp SYBR Q-PCR Mastermix (Thermo Fisher Scientific) and the following primers for each target: GAPDH (5’ GCCTCAAGATCATCAGCAAT 3’) and (5’ CTGTGGTCATGAGTCCTTCC 3’); small tumor antigen: (5’ CAGTGCACAGAAGGCTTTTTGGAACA 3’) and (5’ AGCCTGATTTTGGAACCTGGAGTAGC 3’). The RT-qPCR confirms the previous finding that there is an increase in early protein expression at 48 hours post-infection in the immortalized cells (Figure 1D).

We believe that the RPTE-hTERT cell line will be a useful tool for studying biological aspects of BKPyV infection that cannot be performed in primary cells with limited lifespans in culture.

## Resource availability

RPTE-hTERT cells are provided upon request by contacting the corresponding author.

## Acknowledgment

We thank Eric Campeau & Paul Kaufman for gifting pLenti CMV GFP Puro plasmid through Addgene. This report was supported in part by a Comprehensive Cancer Center Core grnat from the National Cancer Institute of the National Institutes of Health (NIH) under award number P30 CA046592, and NIH grant R01 AI060584 awarded to M.J.I.

